# matchms - processing and similarity evaluation of mass spectrometry data

**DOI:** 10.1101/2020.08.06.239244

**Authors:** Florian Huber, Stefan Verhoeven, Christiaan Meijer, Hanno Spreeuw, Efraín Manuel Villanueva Castilla, Cunliang Geng, Justin J. J. van der Hooft, Simon Rogers, Adam Belloum, Faruk Diblen, Jurriaan H. Spaaks

## Abstract

Mass spectrometry data is at the heart of numerable applications in the biomedical and life sciences. With growing use of high throughput techniques researchers need to analyse larger and more complex datasets. In particular through joint effort in the research community, fragmentation mass spectrometry datasets are growing in size and number. Platforms such as MassBank (Horai et al. 2010), GNPS (Wang et al. 2016) or MetaboLights (Haug et al. 2020) serve as an open-access hub for sharing of raw, processed, or annotated fragmentation mass spectrometry data (MS/MS). Without suitable tools, however, exploitation of such datasets remains overly challenging. In particular, large collected datasets contain data aquired using different instruments and measurement conditions, and can further contain a significant fraction of inconsistent, wrongly labeled, or incorrect metadata (annotations).

---

`Matchms` is an open-access Python package to import, process, clean, and compare mass spectrometry data (MS/MS) (see Figure 1). It allows to implement and run an easy-to-follow, easy-to-reproduce workflow from raw mass spectra to pre- and post-processed spectral data. Raw data can be imported from the commonly used formats msp, mzML, mzXML, MGF (mzML, mzXML, MGF file importers are build on top of pyteomics (Levitsky et al. 2019)(Goloborodko et al. 2013)), as well as from json files (as provided by GNPS), but also via Universal Spectrum Identifiers (USI) (Wang et al. 2020). Further data formats or more extensive options regarding metadata parsing can best be handled by using pyteomics (Levitsky et al. 2019) or pymzml (Kösters et al. 2018). Matchms contains numerous metadata cleaning and harmonizing filter functions that can easily be stacked to construct a desired pipeline (Figure 2), which can also easily be extended by custom functions wherever needed. Available filters include extensive cleaning, correcting, checking of key metadata fields such as compound name, structure annotations (InChI, Smiles, InchiKey), ionmode, adduct, or charge. Many of the provided metadata cleaning filters were designed for handling and improving GNPS-style MGF or json datasets. For future versions, however, we aim to further extend this to other commonly used public databases.

**Figure 1:**
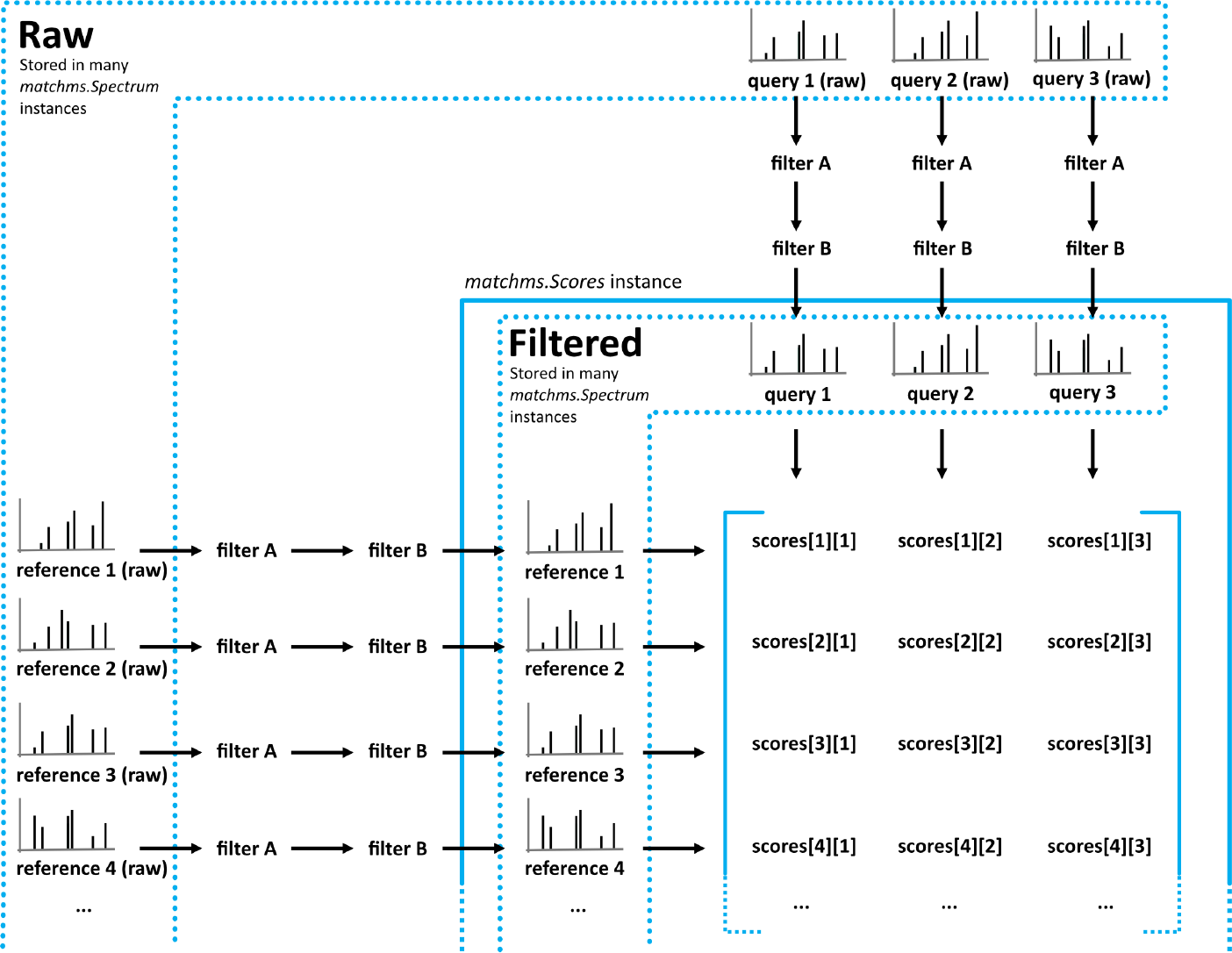
Flowchart of `matchms` workflow. Reference and query spectrums are filtered using the same set of set filters (here: filter A and filter B). Once filtered, every reference spectrum is compared to every query spectrum using the `matchms.Scores` object.

**Figure 2:**
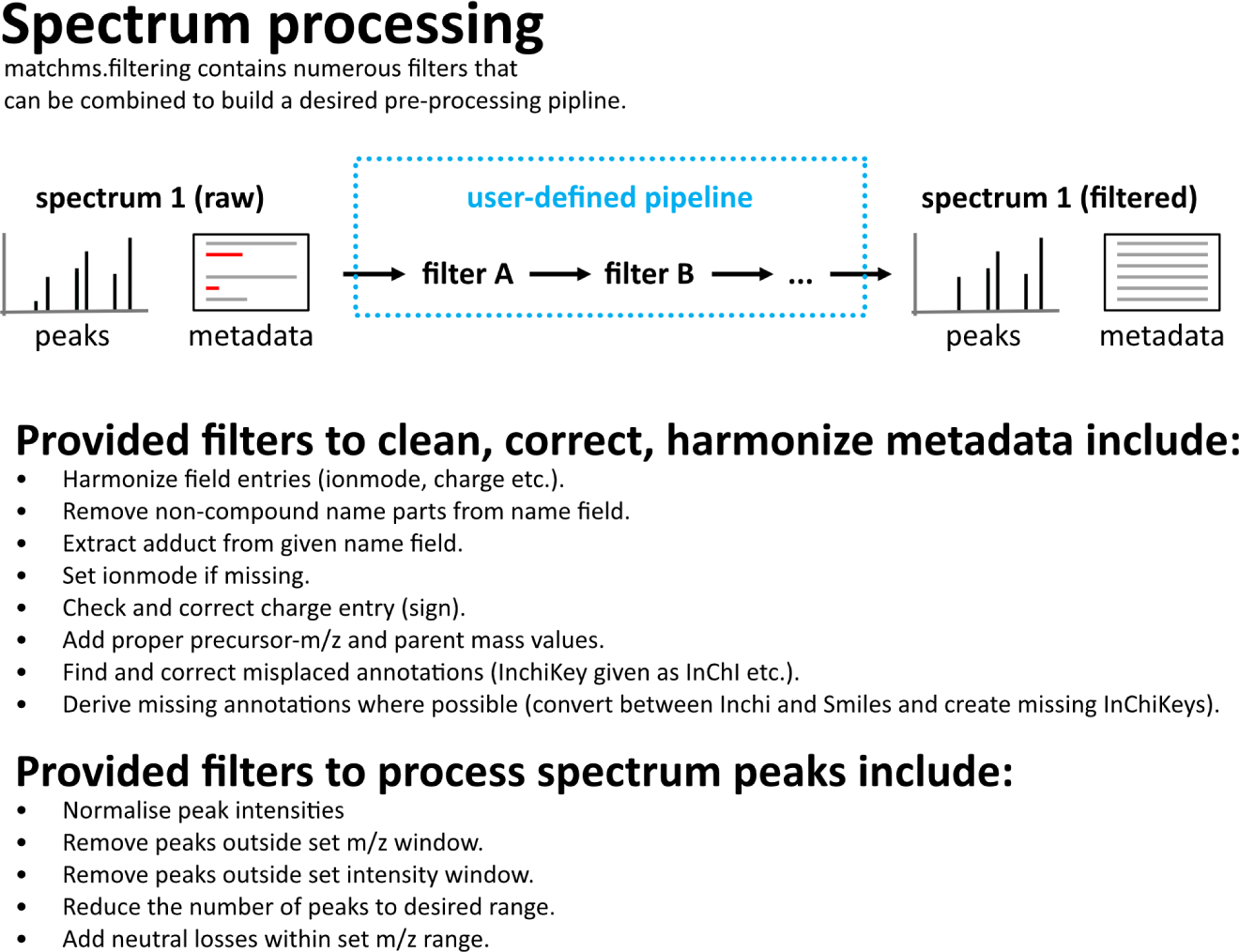
`Matchms` provided a range of filter functions to process spectrum peaks and metadata. Filters can easily be stacked and combined to build a desired pipeline. The API also makes it easy to extend customer pipelines by adding own filter functions.

Current Python tools for working with MS/MS data include pyOpenMS (Röst et al. 2014), a wrapper for OpenMS (Röst et al. 2016) with a strong focus on processing and filtering of raw mass spectral data. pyOpenMS has a wide range of peak processing functions which can be used to further complement a `Matchms` filtering pipeline. Another, more lightweight and native Python package with a focus on spectra visualization is spectrum_utils (Bittremieux 2020). `Matchms` focuses on comparing and linking large number of mass spectra. Many of its build-in filters are aimed at handling large mass spectra datasets from common public data libraries such as GNPS.

`Matchms` provides functions to derive different similarity scores between spectra. Those include the established spectra-based measures of the cosine score or modified cosine score (Watrous et al. 2012). The package also offers fast implementations of common similarity measures (Dice, Jaccard, Cosine) that can be used to compute similarity scores between molecular fingerprints (rdkit, morgan1, morgan2, morgan3, all implemented using rdkit (Landrum, n.d.)). `Matchms` easily facilitates deriving similarity measures between large number of spectra at comparably fast speed due to score implementations based on Numpy (Walt, Colbert, and Varoquaux 2011), Scipy (Virtanen et al. 2020), and Numba (Lam, Pitrou, and Seibert 2015). Additional similarity measures can easily be added using the `matchms` API. The provided API also allows to quickly compare, sort, and inspect query versus reference spectra using either the included similarity scores or added custom measures. The API was designed to be easily extensible so that users can add their own filters for spectra processing, or their own similarity functions for spectral comparisons. The present set of filters and similarity functions was mostly geared towards smaller molecules and natural compounds, but it could easily be extended by functions specific to larger peptides or proteins.

`Matchms` is freely accessible either as conda package (https://anaconda.org/nlesc/matchms), or in form of source-code on GitHub (https://github.com/matchms/matchms). For further code examples and documentation see https://matchms.readthedocs.io/en/latest/. All main functions are covered by tests and continuous integration to offer reliable functionality. We explicitly value future contributions from a mass spectrometry interested community and hope that matchms can serve as a reliable and accessible entry point for handling complex mass spectrometry datasets using Python.

## Example workflow

A typical workflow with matchms looks as indicated in Figure 1, or as described in the following code example.

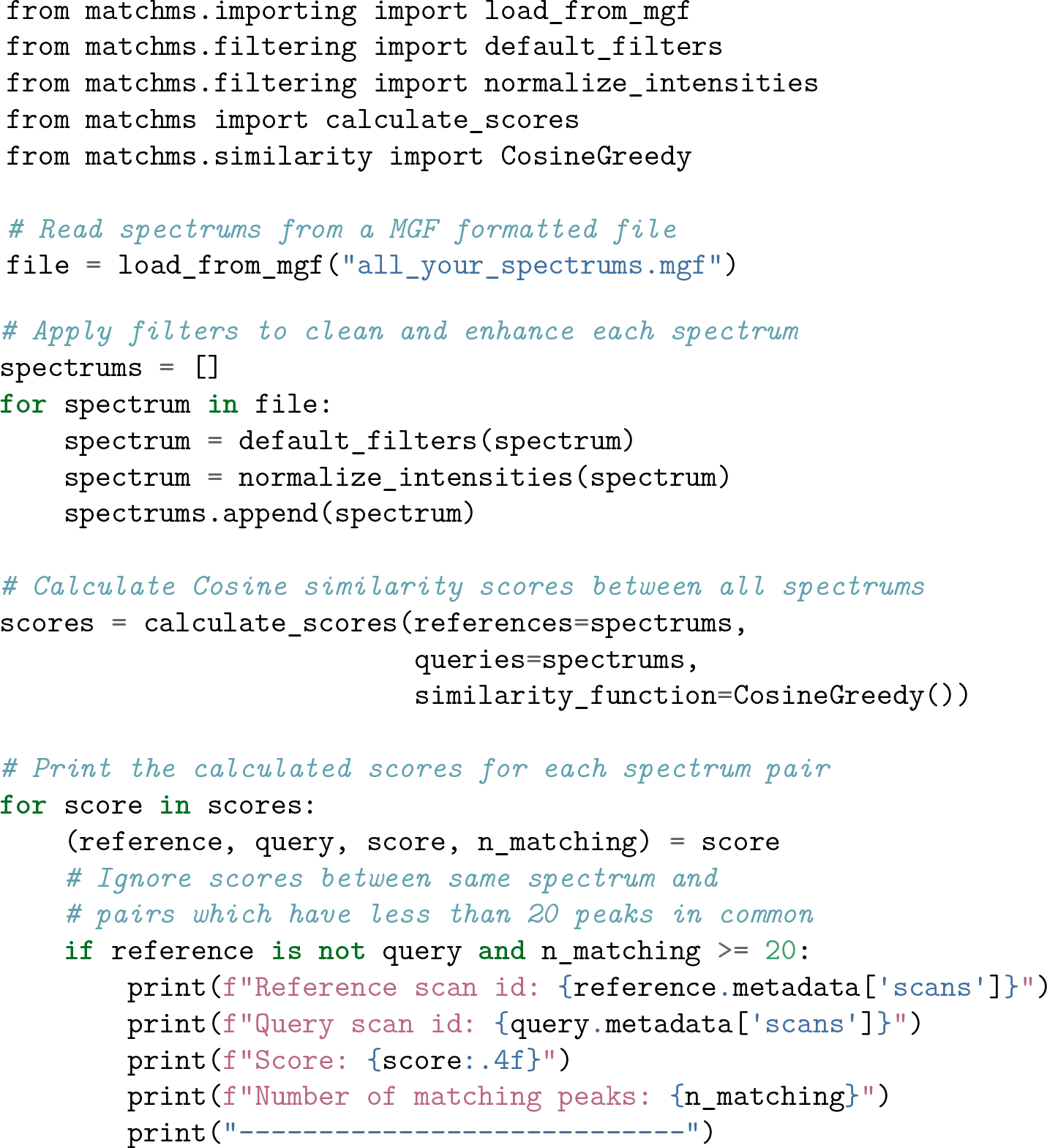

### Processing spectrum peaks and plotting

`Matchms` provides numerous filters to process mass spectra peaks. Below a simple example to remove low intensity peaks from a spectrum (Figure 3).

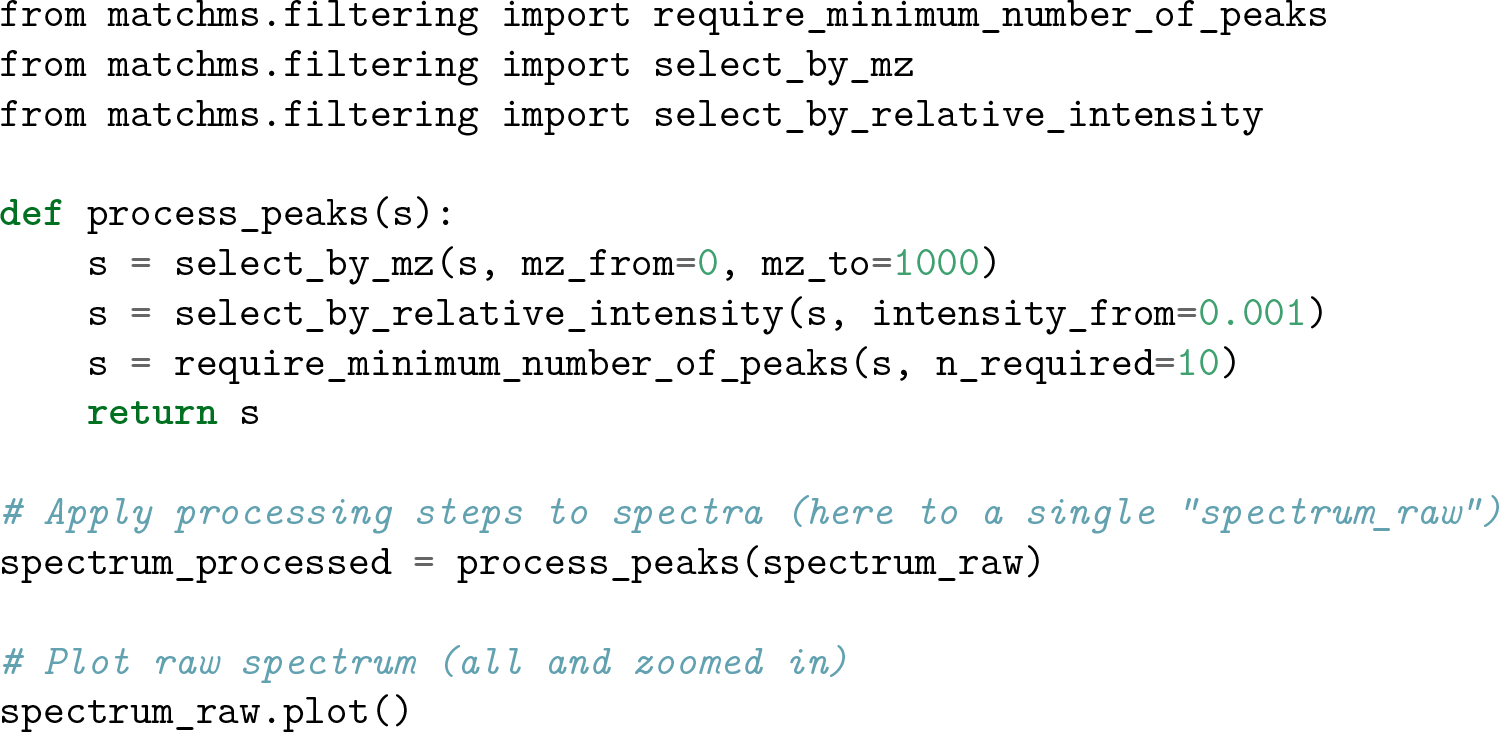

**Figure 3:**
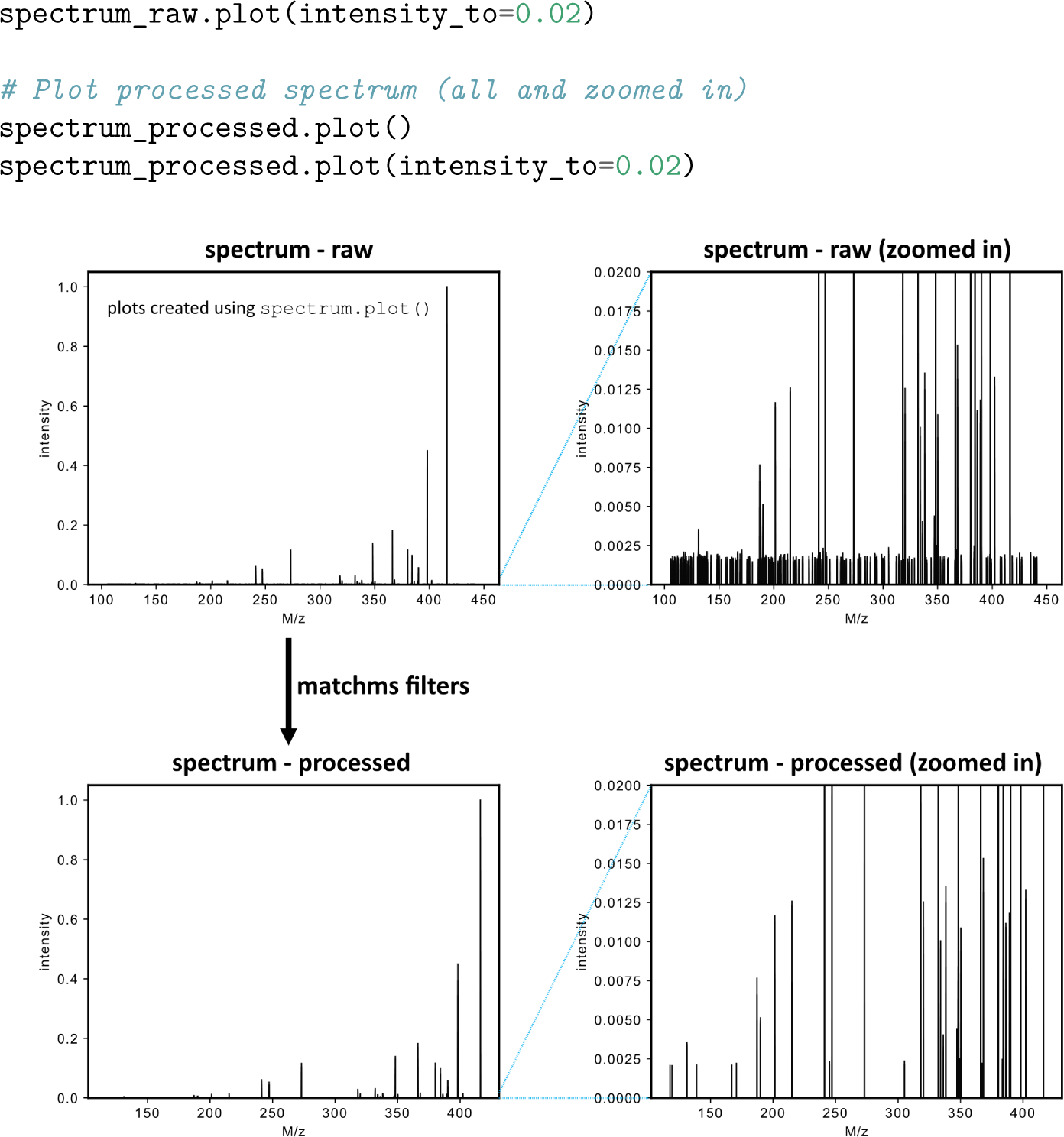
Example of `matchms` peak filtering applied to an actual spectrum using `select_by_relative_intensity` to remove peaks of low relative intensity. Spectra are plotted using the provided `spectrum.plot()` function.

## References

Bittremieux, Wout. 2020. “Spectrum_utils: A Python Package for Mass Spectrometry Data Processing and Visualization.” Analytical Chemistry 92 (1): 659–61. doi:10.1021/acs.analchem.9b04884.

Goloborodko, Anton A., Lev I. Levitsky, Mark V. Ivanov, and Mikhail V. Gorshkov. 2013. “Pyteomics - a Python Framework for Exploratory Data Analysis and Rapid Software Prototyping in Proteomics.” Journal of the American Society for Mass Spectrometry 24 (2): 301–4. doi:10.1021/jasms.8b04453.

Haug, Kenneth, Keeva Cochrane, Venkata Chandrasekhar Nainala, Mark Williams, Jiakang Chang, Kalai Vanii Jayaseelan, and Claire O’Donovan. 2020. “MetaboLights: A Resource Evolving in Response to the Needs of Its Scientific Community.” Nucleic Acids Research 48 (D1): D440–D444. doi:10.1093/nar/gkz1019.

Horai, Hisayuki, Masanori Arita, Shigehiko Kanaya, Yoshito Nihei, Tasuku Ikeda, Kazuhiro Suwa, Yuya Ojima, et al. 2010. “MassBank: A Public Repository for Sharing Mass Spectral Data for Life Sciences.” Journal of Mass Spectrometry 45 (7): 703–14. doi:10.1002/jms.1777.

Kösters, M., J. Leufken, S. Schulze, K. Sugimoto, J. Klein, R. P. Zahedi, M. Hippler, S. A. Leidel, and C. Fufezan. 2018. “pymzML V2.0: Introducing a Highly Compressed and Seekable Gzip Format.” Bioinformatics 34 (14): 2513–4. doi:10.1093/bioinformatics/bty046.

Lam, Siu Kwan, Antoine Pitrou, and Stanley Seibert. 2015. “Numba: A LLVM-Based Python JIT Compiler.” In Proceedings of the Second Workshop on the LLVM Compiler Infrastructure in HPC, 1–6. LLVM ‘15. Austin, Texas: Association for Computing Machinery. doi:10.1145/2833157.2833162.

Landrum, Greg. n.d. “RDKit: Open-Source Cheminformatics.” http://www.rdkit.org.

Levitsky, Lev I., Joshua A. Klein, Mark V. Ivanov, and Mikhail V. Gorshkov. 2019. “Pyteomics 4.0: Five Years of Development of a Python Proteomics Framework.” Journal of Proteome Research 18 (2): 709–14. doi:10.1021/acs.jproteome.8b00717.

Röst, Hannes L., Timo Sachsenberg, Stephan Aiche, Chris Bielow, Hendrik Weisser, Fabian Aicheler, Sandro Andreotti, et al. 2016. “OpenMS: A Flexible Open-Source Software Platform for Mass Spectrometry Data Analysis.” Nature Methods 13 (9): 741–48. doi:10.1038/nmeth.3959.

Röst, Hannes L., Uwe Schmitt, Ruedi Aebersold, and Lars Malmström. 2014. “pyOpenMS: A Python-Based Interface to the OpenMS Mass-Spectrometry Algorithm Library.” PROTEOMICS 14 (1): 74–77. doi:10.1002/pmic.201300246.

Virtanen, Pauli, Ralf Gommers, Travis E. Oliphant, Matt Haberland, Tyler Reddy, David Cournapeau, Evgeni Burovski, et al. 2020. “SciPy 1.0: Fundamental Algorithms for Scientific Computing in Python.” Nature Methods 17: 261–72. doi:https://doi.org/10.1038/s41592-019-0686-2.

Walt, Stefan van der, S. Chris Colbert, and Gael Varoquaux. 2011. “The NumPy Array: A Structure for Efficient Numerical Computation.” Computing in Science Engineering 13 (2): 22–30. doi:10.1109/MCSE.2011.37.

Wang, Mingxun, Jeremy J. Carver, Vanessa V. Phelan, Laura M. Sanchez, Neha Garg, Yao Peng, Don Duy Nguyen, et al. 2016. “Sharing and Community Curation of Mass Spectrometry Data with Global Natural Products Social Molecular Networking.” Nature Biotechnology 34 (8): 828–37. doi:10.1038/nbt.3597.

Wang, Mingxun, Simon Rogers, Wout Bittremieux, Christopher Chen, Pieter C. Dorrestein, Emma L. Schymanski, Tobias Schulze, Steffen Neumann, and Rene Meier. 2020. “Interactive MS/MS Visualization with the Metabolomics Spectrum Resolver Web Service.” bioRxiv, May, 2020.05.09.086066. doi:10.1101/2020.05.09.086066.

Watrous, Jeramie, Patrick Roach, Theodore Alexandrov, Brandi S. Heath, Jane Y. Yang, Roland D. Kersten, Menno van der Voort, et al. 2012. “Mass Spectral Molecular Networking of Living Microbial Colonies.” Proceedings of the National Academy of Sciences of the United States of America 109 (26): E1743–1752. doi:10.1073/pnas.1203689109.

